# Synthesis and modification of a mesoporous material type MCM-41 by an amine for the adsorption of organic pollutants: anionic and cationic dyes

**DOI:** 10.1101/118182

**Authors:** Benyoub Nassima, Benhamou Abdallah, Debab Abdelkader

**Affiliations:** Departement of Chemical Engineering, Faculty of Chemistry, University of Scinece and Technology MOHAMED BOUDIAF –ORAN, El Mnaouar, BP 1505, Bir El Djir 31000

**Keywords:** Mesoporous materials, MCM-41, adsorption, Orange II Sodium, Janus Green B, adsorption of dyes

## Abstract

The main objective of this work was to synthesize the MCM-41 material from optimized protocols. In the second step, the pore size and the specific surface area of the parent material was increased by the incorporation of swelling agents (long chain carbon amines N-N, Dimethyl-dodecylamine « DMDDA ») in post-synthesis. Then the selective extraction of the amine and the calcination allowed us to obtain materials with pore sizes and even larger surfaces than those of the starting materials (parent materials). The adsorbents, identified as MCM-41 (P), DMDDA-41 (A), DMDDA-41 (B), DMDDA-41 (C) and MCM-41 (P/C), were characterized regarding their texture, mesoscopic ordering and chemical surface and finally their adsorption capacity was evaluated using two diffrent dyes (anionic: Orange II sodium and cationic: Janus Green B). The results obtained during the adsorption study show the efficiency of these materials, in particular the "amine and calcined" materials for decolorizing aqueous media contaminated with organic dyes. The kinetic studies and the adsorption isotherms were carried out to clarify the method of fixing each of the two dyes on the two materials tested. The experiments showed that the amine material had a maximum capacity for fixing Orange II (221.06 mg / g) (anionic dye), whereas for a maximum adsorption capacity (455.23 mg / g) Of the Janus GB (cationic dye), the calcined material is more efficient.

## 1 Introduction

Since the 1990s, the chemistry of inorganic nanostructured materials has developed considerably with the introduction of soft sol-gel chemistry. Thus, the scientists of the Mobil company proposed the first syntheses of mesostructured silicates, ie materials with an organized porous system made up of mesopores [1, 2]. Since then, many research groups have patented new families of materials with different pore structures, pore sizes and synthesis methods: several methods are needed in the manufacturing process of new porous materials organized "MPO".

At present, a new family of ordered mesoporous solids is very widely studied by many researchers from different horizons for various applications including adsorption [3, 4] and catalysis. In the field of adsorption, mesoporous materials such as MCM-41, HMS, SBA-15, SBA-1 have been functionalized by various groups for the adsorption of metal ions and various organic pollutants (synthetic dyes) [5]. The main objective of this work is to synthesize a mesoporous material of type MCM-41 (hexagonal), to modify it by: addition of amine, selective extraction with ethanol and calcination and then d To evaluate the adsorption capacity of these mesoporous materials with respect to two dyes, anionic sodium orange II and Janus Green B with a cationic character. A detailed study is presented defining the different methods of characterization, as well as a kinetic study and that of adsorption isotherms will also be discussed.L’objectif principal de ce

## 2 Synthesis and Characterization

### 2.1. Materials

The principal reagents used are summarized in a table which follows, where their origin, formula and possibly their purity are specified (Table 1).

**Table 1:**
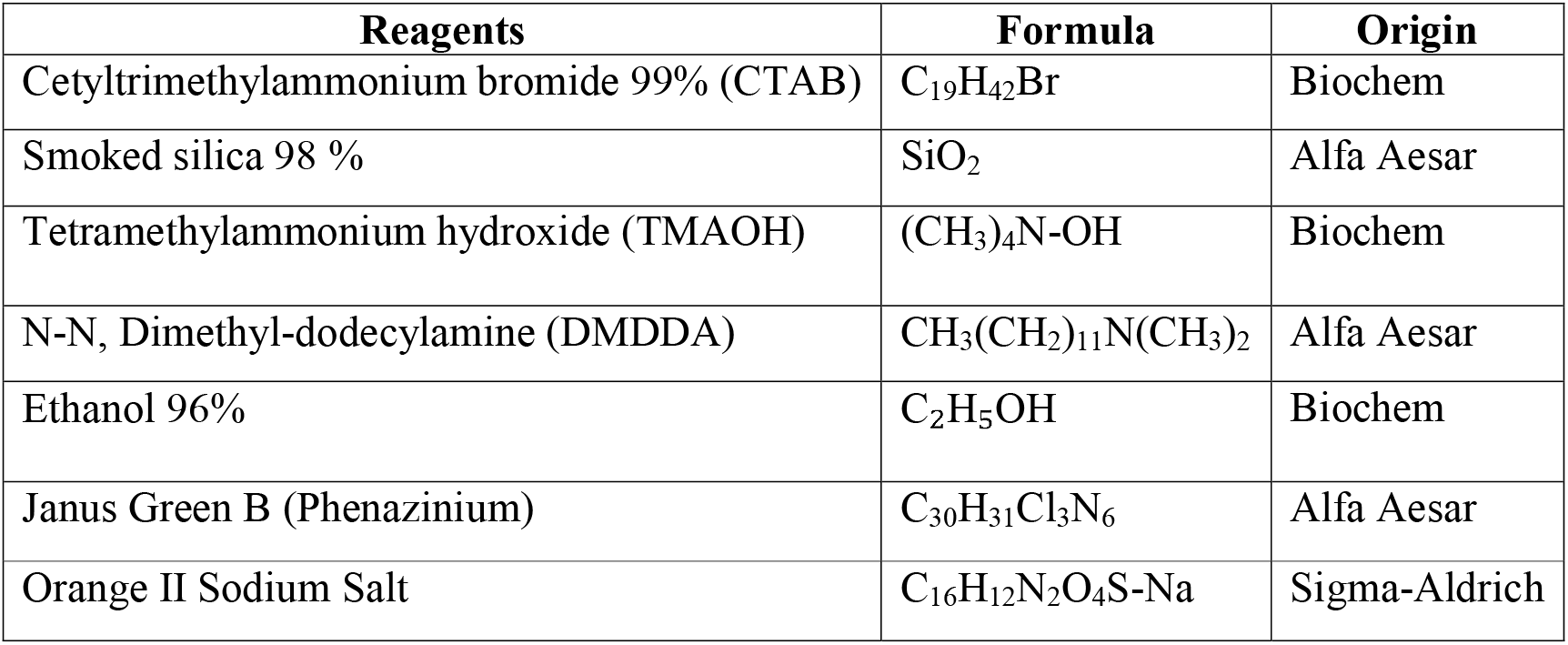
Main reagents used.

| Reagents | Formula | Origin |
| --- | --- | --- |
| Cetyltrimethylammonium bromide 99% (CTAB) | $\text{C}_{19}\text{H}_{42}\text{Br}$ | Biochem |
| Smoked silica 98 % | $\text{SiO}_2$ | Alfa Aesar |
| Tetramethylammonium hydroxide (TMAOH) | $(\text{CH}_3)_4\text{N-OH}$ | Biochem |
| N-N, Dimethyl-dodecylamine (DMDDA) | $\text{CH}_3(\text{CH}_2)_{11}\text{N}(\text{CH}_3)_2$ | Alfa Aesar |
| Ethanol 96% | $\text{C}_2\text{H}_5\text{OH}$ | Biochem |
| Janus Green B (Phenazinium) | $\text{C}_{30}\text{H}_{31}\text{Cl}_3\text{N}_6$ | Alfa Aesar |
| Orange II Sodium Salt | $\text{C}_{16}\text{H}_{12}\text{N}_2\text{O}_4\text{S-Na}$ | Sigma-Aldrich |

In all cases, the solvent employed is distilled water and, the syntheses are carried out at ambient temperature, in an open container. All are made with stirring.

### 2.2. S ynthesis

In a beaker, the water / TMAOH mixture is stirred vigorously for 5 minutes, then the CTAB is added in small quantities without stopping the stirring which will continue for 30 minutes, and finally the SiO.sub.2 is added in order to have A gel after about 2 hours (the speed of agitation is less strong).

The mixture is transferred to the Teflon reactor and placed in an oven set at 90 ° to 100 ° C. for 48 hours. Afterwards, one part of the surfactant is removed by Büchner filtration (filtration and washing with distilled water) and then placed in an oven for drying for one day at 100 ° C. The material obtained has the following molar composition: 1 SiO_2_; 0.25 CTAB; 0.2 TMAOH; 40 H_2_O [6, 7], is named: parent material « Si-MCM-41-P ».

Si-MCM-41, called material P, is added to an emulsion consisting of water and N-N-Dimethyl-dodecylamine (DMDDA) previously stirred for 5 minutes. This mixture is left under stirring for 30 minutes, then transferred to a Teflon reactor, which is placed in an oven set at 120°C. for 3 days. The material obtained is filtered and washed several times with distilled water and then dried at ambient temperature: it is the « **Si-MCM-41/A** » amine material.

The selective extraction of the amine is carried out using a specific solvent which is the ethanol, using a soxhlet. The material obtained is the « **Si-MCM-41/B** » deaminated material.

The material Si-MCM-41 / P is calcined in 550°C during 7 hours giving rise to a material called « **Si-MCM-41-P / C** » parent / calcined material.

This step is carried out at 550°C. for 7 hours with a heating rate of 1°C./min, giving rise to a material called « **Si-MCM-41/C** » calcined material.

### 2.3. Characterization techniques

This characterization aims in a very thorough way, the study and the identification of the textural properties, structural and superficial, as well as the electrochemical properties of the surface of the obtained materials.

A combination of physicochemical (X-ray diffraction, BET, ATG, ATD and Zetrametry) and spectroscopic (FTIR) techniques is realized and their understandings could allow a good exploitation of these materials for specific applications.

## 3 Results & discussion

The X-ray diffraction at low angles is used to demonstrate the arrangement of the channels created by the micelles of surfactants.

Figure 1 shows the diffractograms measured between 0.5 and 6 degrees (2θ) of the DMDDA-41A, DMDDA-41B and DMDDA-41C materials, respectively the DMDDA amine material, the previous material after deamination, and Calcined material. These four materials are compared to the MCM-41/P which is the parent material.

**Figure 1:**
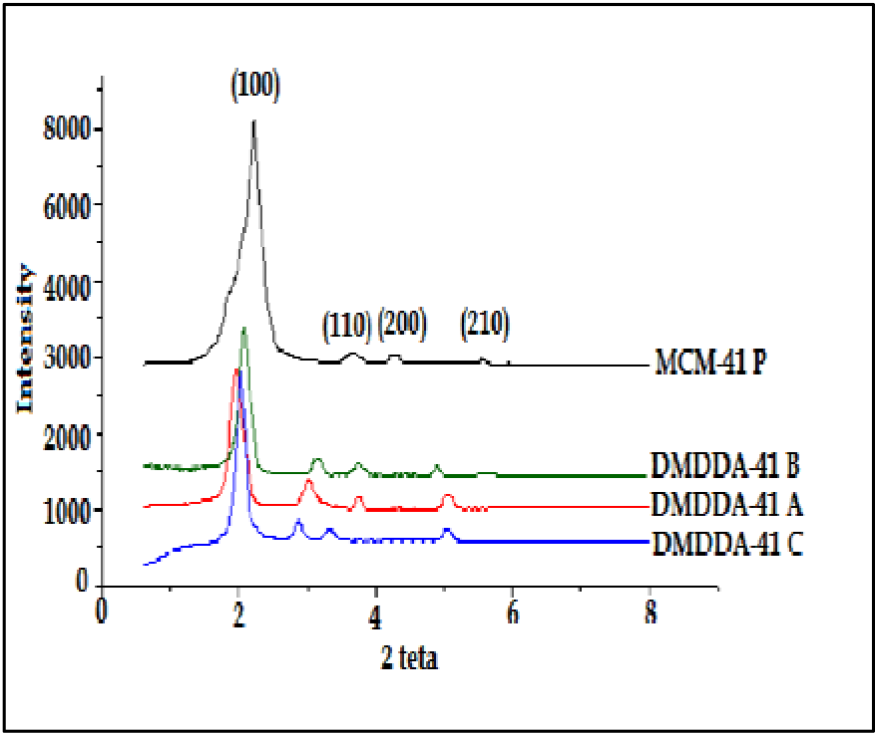
Si-MCM-41 diffraction pattern (parent, amine, deaminated, calcined)

The main peak is observed at 2θ = 2.28° followed by three less intense peaks between 3.2 <2θ <5.4°. This figure can be indexed in a crystallographic system and it is possible to attribute the peaks to diffractions on reticular planes indexed by Miller indices "hkl": the planes (100), (110), (200) and 210) as shown in the figure.

The diffractograms of the materials (amine, deaminated and calcined) have the same peaks as those observed on the purely silicic material MCM-41/P, which shows that the structure of the material is preserved after modifications made more than they all present an arrangement 2D hexagonal (p6mm) of the pores (Figure 2). [8]

**Figure 2:**
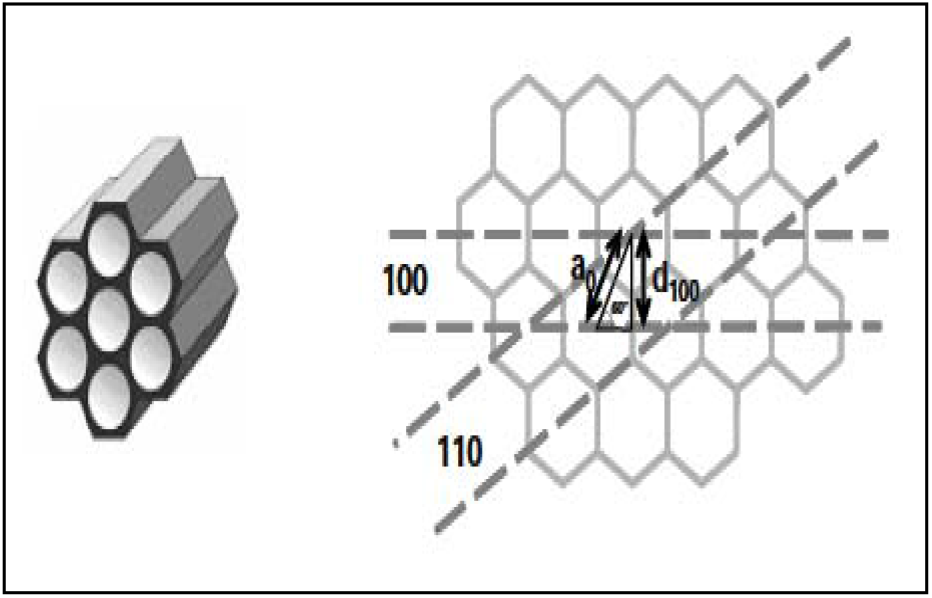
Diagram of the pore arrangement.

A peak slippage (MCM-41P and DMDDA-41B) is observed at higher diffraction angles [9, 10, 13], which results in smaller inter-reticular distances as shown in the table.

Table 2 summarizes the structural data derived from the diffractograms for all materials prepared from MCM-41.

**Table 2:**
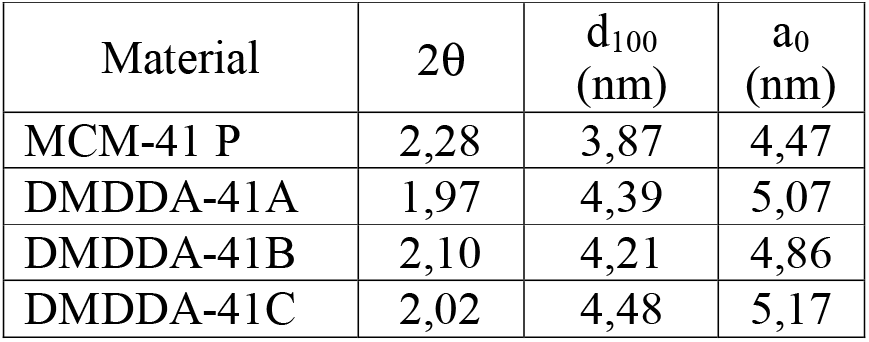
X-ray diffraction data of MCM-41P and modified materials.

The adsorption-desorption isotherms of the parent MCM-41 and the prepared materials are shown in figure 3. These are type IV according to the IUPAC classification and Hysteresis H1 which is representative of structured mesopores. [11, 12, 13, 30]

**Figure 3:**
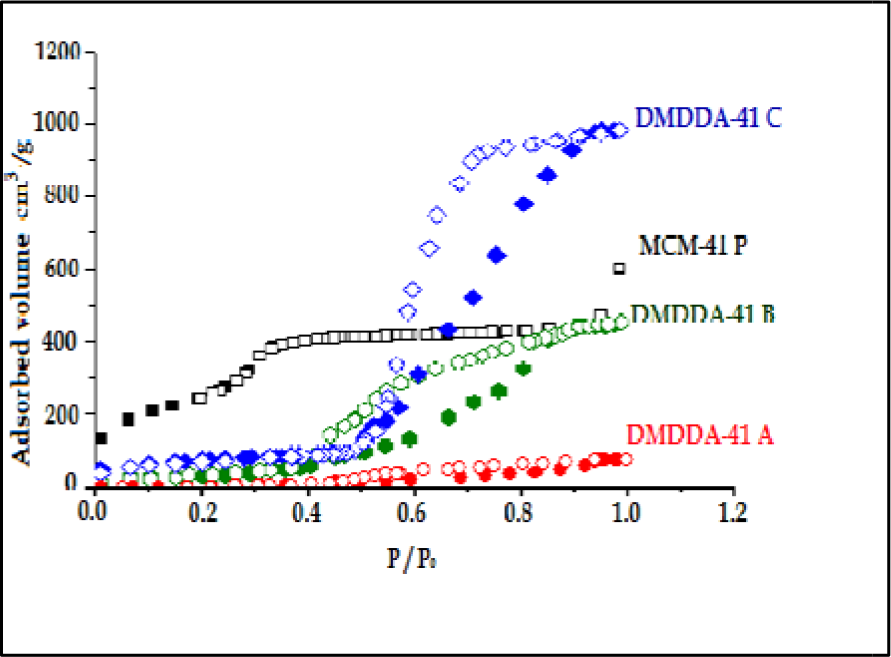
Isotherm of adsorption / desorption of Si-MCM-41.

For the MCM-41P, an increasing slope is observed for low relative pressures 0 <(P / P0) <0.3 corresponding to a monolayer filling of the surface [10], followed by a steep slope of the curve for the Relative pressures between 0.3 <(P / P0) <0.8; Due to the capillary condensation of the nitrogen inside the mesopores. And towards (P / P0)> 0.8 there is a multilayer adsorption at the surface of the parent material.

For the DMDDA-41A material, at relative pressures P / P0 <0.4, a monolayer filling of the surface is obtained and for P / P0> 0.5 there appears a plateau of a multilayer adsorption. The same observations noted for the parent material are noted for DMDDA-41B and DMDDA-41C. For both materials, the relative low pressure plateau is almost the same (P / P0 <0.5), but the distinction is in the slopes of the following plateau: steep slope to the calcined material (up to 1 in P / P0) in contrast to the deaminated material (0.5 <P / P0 <0.8).

In the case of these two materials, the desorption does not follow the adsorption creating an H1 type hysteresis which abruptly closes at P / P0 = 0.5 for the material (DMDDA-41C) and at P / P0 = 0.4 For the DMDDA-41B material, this suggests a phenomenon of capillary condensation in the mesopores. The width of the hysteresis increases (DMDDA-41C) indicating the pore size distribution is much wider in the calcined material (Figure 4) [13, 23, 24].

**Figure 4:**
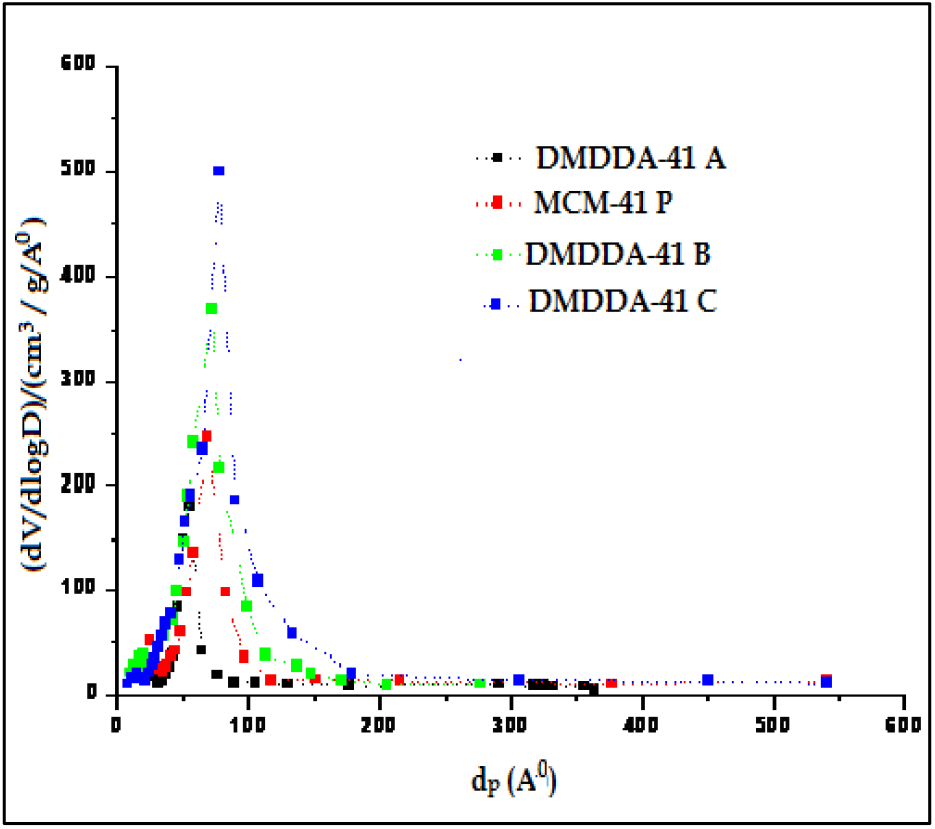
Distribution of pore diameters of materials MCM-41.

The pore diameter distributions shown in figure 4 show that the materials prepared have different pore diameters. The traces are all centered indicating a better uniformity of the pore diameter. As the pore diameter distribution widens, this indicates the formation of a less homogeneous network of pores.

The analysis of these data makes it possible to observe great differences in the parameters of the four materials. The results of surface area, pore diameter and pore volume decrease remarkably for the amine material (the surface and part of the pore volume are occupied by the amines), in contrast to MCM-41P.

Afterwards, deamination increases these parameters again, while calcination leads to higher values.

**Table 3:** Comparison of nitrogen adsorption-desorption data at 77 ° K of MCM-41P and modified materials.

| Material | $S_{BET}$<br>($m^2/g$) | $d_p$<br>(nm) | $V_p$<br>($cm^3/g$) |
| --- | --- | --- | --- |
| MCM-41 P | 1147 | 3,22 | 0,85 |
| DMDDA-41A | 74 | - | 0,28 |
| DMDDA-41B | 387 | 5,84 | 1,07 |
| DMDDA-41C | 1246 | 10,6 | 1,86 |

The IRTF spectra of the MCM-41 materials prepared are shown in figure 5 where it is found that some vibration bands are present in the materials: they are similar to that of amorphous silica.

**Figure 5:**
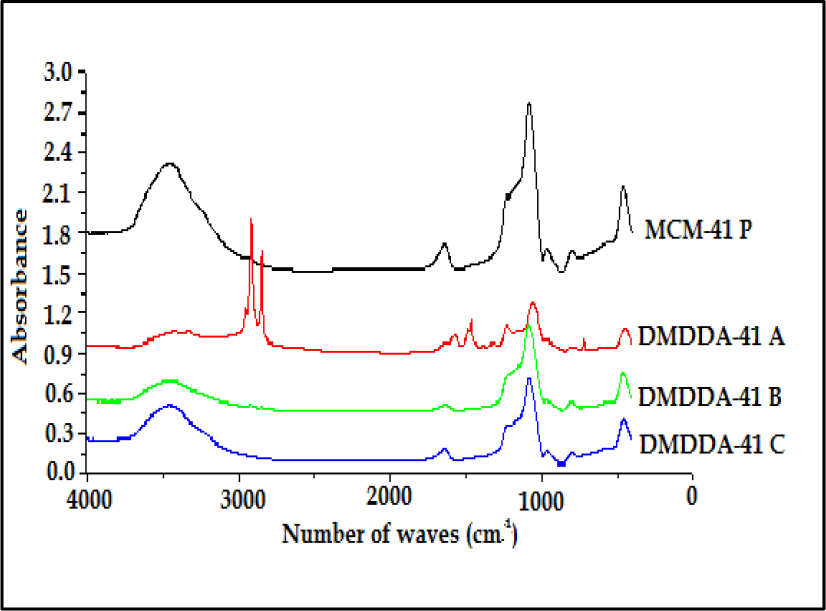
IRTF Spectrum of Si-MCM-41.

The broad band between 3450 cm^−1^, characteristic of elongation of the bond (OH), water and silanol groups of the surface. Another vibrating band (O-H) noted around 1650 cm^−1^ testifying to the water present in the materials. At 1100 cm^−1^, an asymmetric (O-Si-O) elongation junction of the SiO_4_ tetrahedral entities was observed, whereas about 950 and 750 cm^−1^, two bends characteristic of the asymmetric elongation of the (Si-O) bond of these same entities present. We note also the presence of a band characterizing the deformation of the angle (O-Si-O) of the tetrahedral entities SiO4, this towards 450cm^−1^. [14, 16, 21]

In addition to these bands, the amine material (DMDDA-41A) records two new bands characteristic of the amine groups, the first located at 2900 cm−1 characterizes the vibrations (CN), while the second one is observed at 1480 cm −1 Resulting from the deformation vibration of the bonds (NH). [14, 18, 22]

The ATG / ATD thermogravimetric analysis of all the materials is represented by two figure6 (A and B), from which there are three different zones of mass variation as a function of temperature.

**Figure 6:**
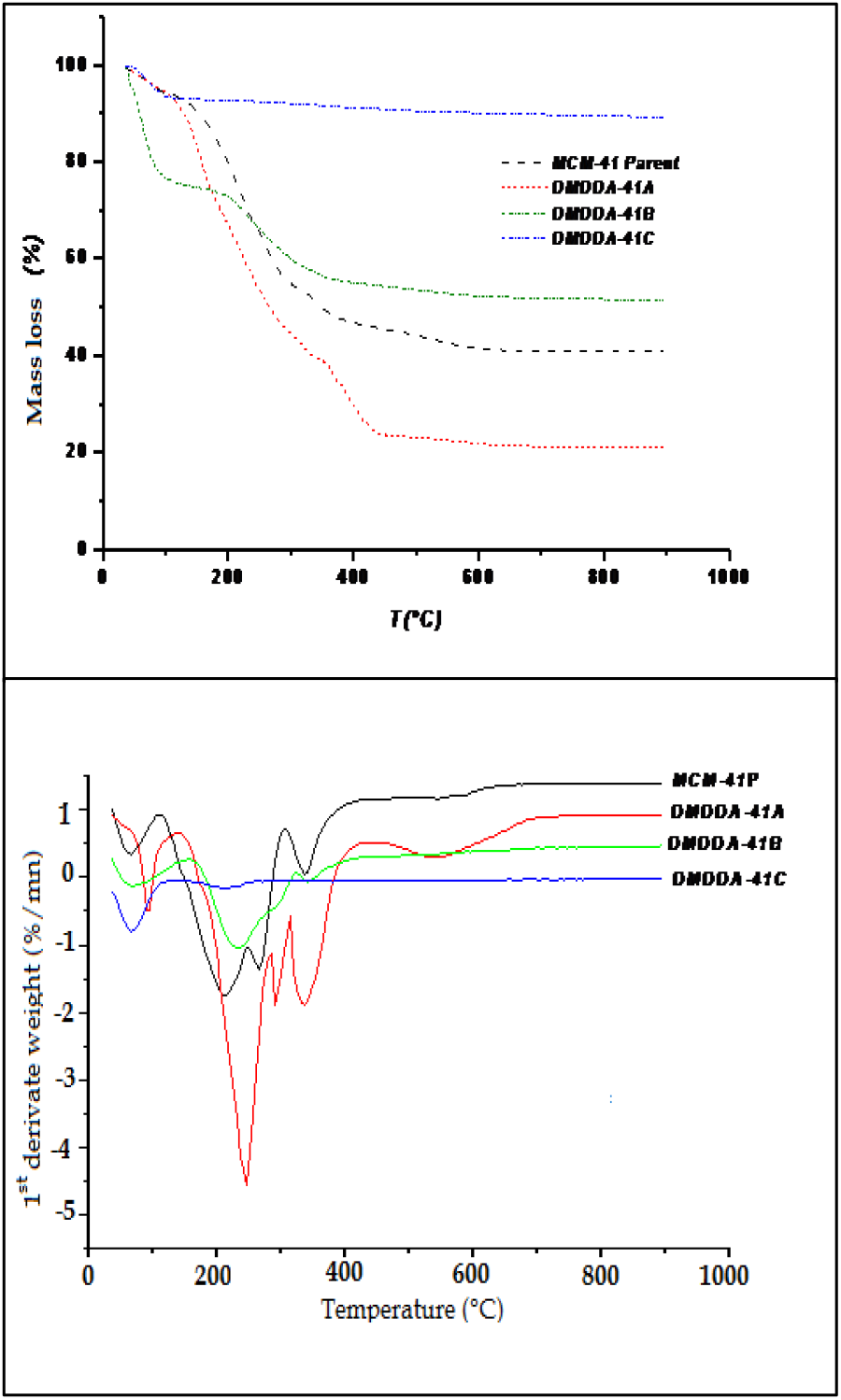
Thermogravimetric analysis A-(TGA) & Differential thermal analysis B-(DTA) of the different materials.

The 1^st^ zone at low temperature (25<T <160°C.) corresponds to dehydration of the water physisorbed on the surface of the material. The 2^nd^ intermediate zone (160<T <600°C) corresponds to the decomposition and volatilization of the organic compounds (surfactants and amine) in strong interaction with the surface of the material (covalent and electrostatic bonds) The last high temperature zone (600<T <900°C):assigned to silica dehydroxylation phenomena (condensation of the remaining silanols causing the elimination of water molecules) [8, 10, 15, 17, 24].

**Table 4:** Thermogravimetric (TGA) and Thermal Differential (ATD) data for the different materials.

| Sample | Mass loss (%) |  |  |
| --- | --- | --- | --- |
|  | 1<sup>st</sup> Zone | 2<sup>nd</sup> Zone | 3<sup>rd</sup> Zone |
| MCM-41 P | 8,23 | 48,41 | 1,43 |
| DMDDA -41 A | 3,51 | 70,67 | 3,59 |
| DMDDA -41 B | 15,67 | 45,97 | 1,63 |
| DMDDA -41 C | 7,02 | 2,76 | 0,87 |

Figure 7 shows the curves ζ = f (pH) of the various MCM-41 materials in the pH range from 2 to 11. Each point ζ = f (pH) is the mean of the results obtained on the measurements of the electrophoretic mobility of 100 to 500 p at a zeta potential of between 5 and 8 mV. The pH for which the zeta potential is zero (no displacement of the particles under the effect of the electric field) is called the isoelectric point (PIE). [19, 20]

Parent material MCM-41 has positive potentials between 2 <pH <5 and negative potentials from pH = 5.5. For the DMDDA-41 A, it has positive potentials between 2 <pH <8.5 and negative potentials from pH = 8.5. Then, DMDDA-41 B has positive potentials between 2 <pH <6 and negative potentials from pH = 6.1.

And for the calcined material, it has positive potentials between 2 <pH <3.6 and negative potentials from pH = 3.6 [16, 18, 23, 24].

**Table 5:**
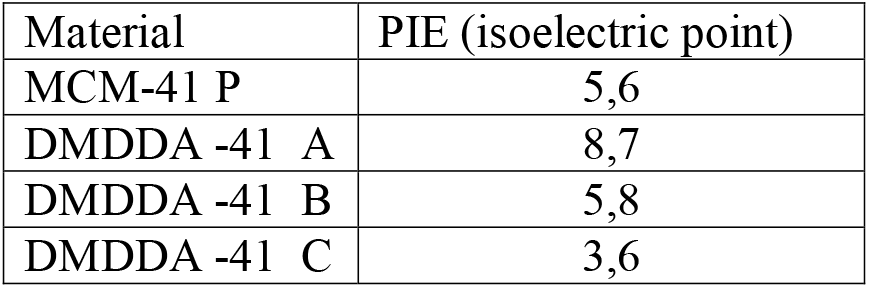
Values of the isoelectric points of the different materials.

| Material | PIE (isoelectric point) |
| --- | --- |
| MCM-41 P | 5,6 |
| DMDDA -41 A | 8,7 |
| DMDDA -41 B | 5,8 |
| DMDDA -41 C | 3,6 |

## 4 Application to adsorption

Our study consists in developing techniques for low - cost water discolouration while recovering industrial waste: Discoloration of synthetic solutions: ***Orange 2 sodium*** with anionic character [25] and ***Janus Green B*** with a cationic character. Their characteristic data are summarized in the following table.

**Table 6:**
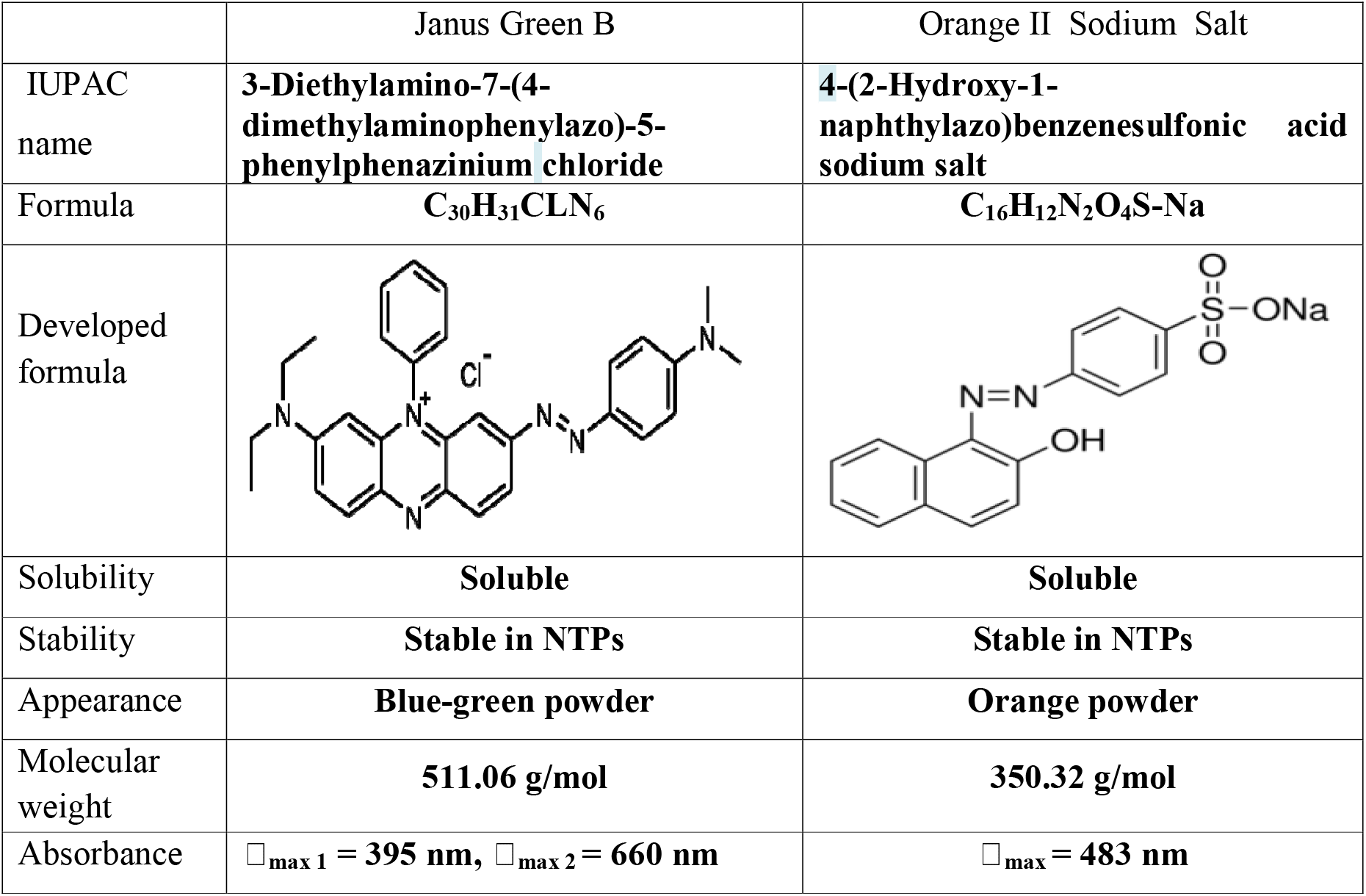
Physico-chimical characteristics of the two dyes.

|  | Janus Green B | Orange II Sodium Salt |
| --- | --- | --- |
| IUPAC name | **3-Diethylamino-7-(4-dimethylaminophenylazo)-5-phenylphenazinium chloride** | **4-(2-Hydroxy-1-naphthylazo)benzenesulfonic acid sodium salt** |
| Formula | **C<sub>30</sub>H<sub>31</sub>CLN<sub>6</sub>** | **C<sub>16</sub>H<sub>12</sub>N<sub>2</sub>O<sub>4</sub>S-Na** |
| Developed formula |  |  |
| Solubility | **Soluble** | **Soluble** |
| Stability | **Stable in NTPs** | **Stable in NTPs** |
| Appearance | **Blue-green powder** | **Orange powder** |
| Molecular weight | **511.06 g/mol** | **350.32 g/mol** |
| Absorbance | $\lambda_{\text{max } 1} = 395 \text{ nm}, \lambda_{\text{max } 2} = 660 \text{ nm}$ | $\lambda_{\text{max}} = 483 \text{ nm}$ |

First, a study of their UV-visible spectra at wavelengths between 200 and 800 nm was carried out to determine the wavelengths which correspond to the maximum adsorption and to establish the calibration curves. Thus the 483 nm, 618 nm waves were respectively for the Orange II Sodium Salt and the Janus Green B. We then studied the influence of different physicochemical parameters on the coloring interactions – synthesized materials (agitation time, Initial dye concentration, adsorption capacity test).

It is noted that in each experiment, the tests were carried out in flasks covered with aluminum foil, in order to avoid degradation of the pollutant in the presence of light.

The study of the adsorption of the two dyes on the different materials is represented in the histograms of the quantities adsorbed by the various materials synthesized (figure 8)

**Figure 8:**
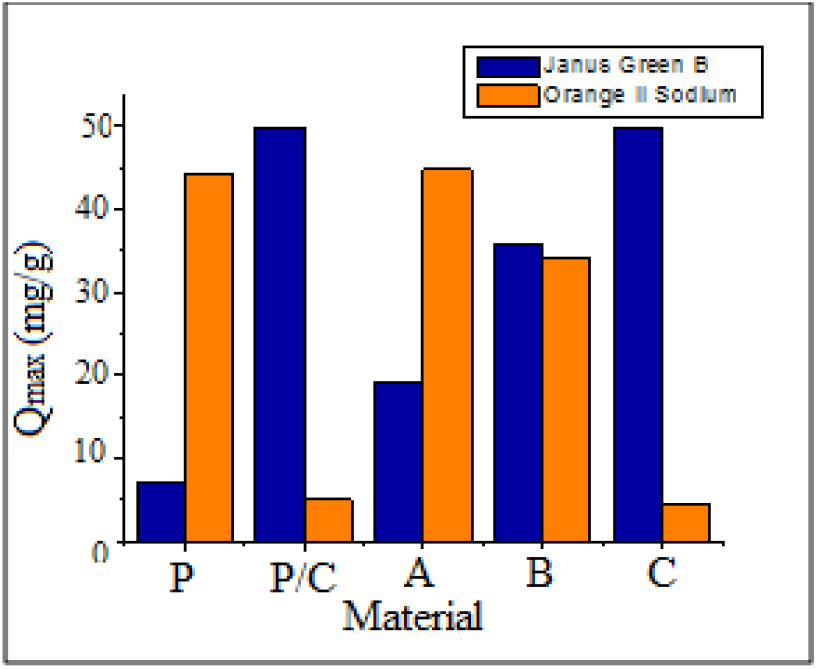
The retention quantities of the two dyes by the various materials MCM-41.

We note that the two materials "parent and amine" possess the greatest adsorption capacity of the anionic dye "Orange II". On the other hand, the two materials, namely the calcined parent and the calcined modifier, possess the highest adsorption rates of the cationic dye "Janus GB". The DMDDA-41 B exhibits in both dyes an average adsorption

This difference in the adsorption rates of the different materials is due to two reasons:

1. Differences in the physico-chemical properties of materials (porosity, S_BET_, …)
2. Differences in surface loads of different materials.

The parent material has only the silanol (Si-OH) groups and quaternary ammonium groups of the surfactant, which explains its ability to fix the anionic dye more than the cationic. Moreover, the amine material has, in addition to the preceding groups [27], amine groups (NH.sub.2): the charge of their surface is more positive at the maximum capacity for fixing the Orange II.

For calcined materials (DMDDA-41 P / C and DMDDA-41 C), they have only the silanol groups (Si-OH) which make the charge of their surface more negative, which makes their adsorption capacity of the Janus GB maximum.

**Table 7:** Retention quantities of Orange II Sodium and Janus Green B by the different materials.

| Material<br>MCM-41 | P | P/C | A | B | C |
| --- | --- | --- | --- | --- | --- |
| $Q_{\max}$ (mg/g)<br>« Orange II Sodium » | 44.34 | 4.96 | 44.80 | 33.98 | 4.28 |
| $Q_{\max}$ (mg/g)<br>« Janus GB » | 7.05 | 49.80 | 19.13 | 35.70 | 49.90 |

The adsorption kinetics tests are carried out on the amine and calcined matrix, since they possess, respectively, the greatest adsorption capacity of the anionic and cationic micropollutant. The results are presented by the evolution of the adsorbed quantities of the two dyes by the two materials (Qe) as a function of the contact time (t) in FIG. 9 (A and B).

**Figure 9:**
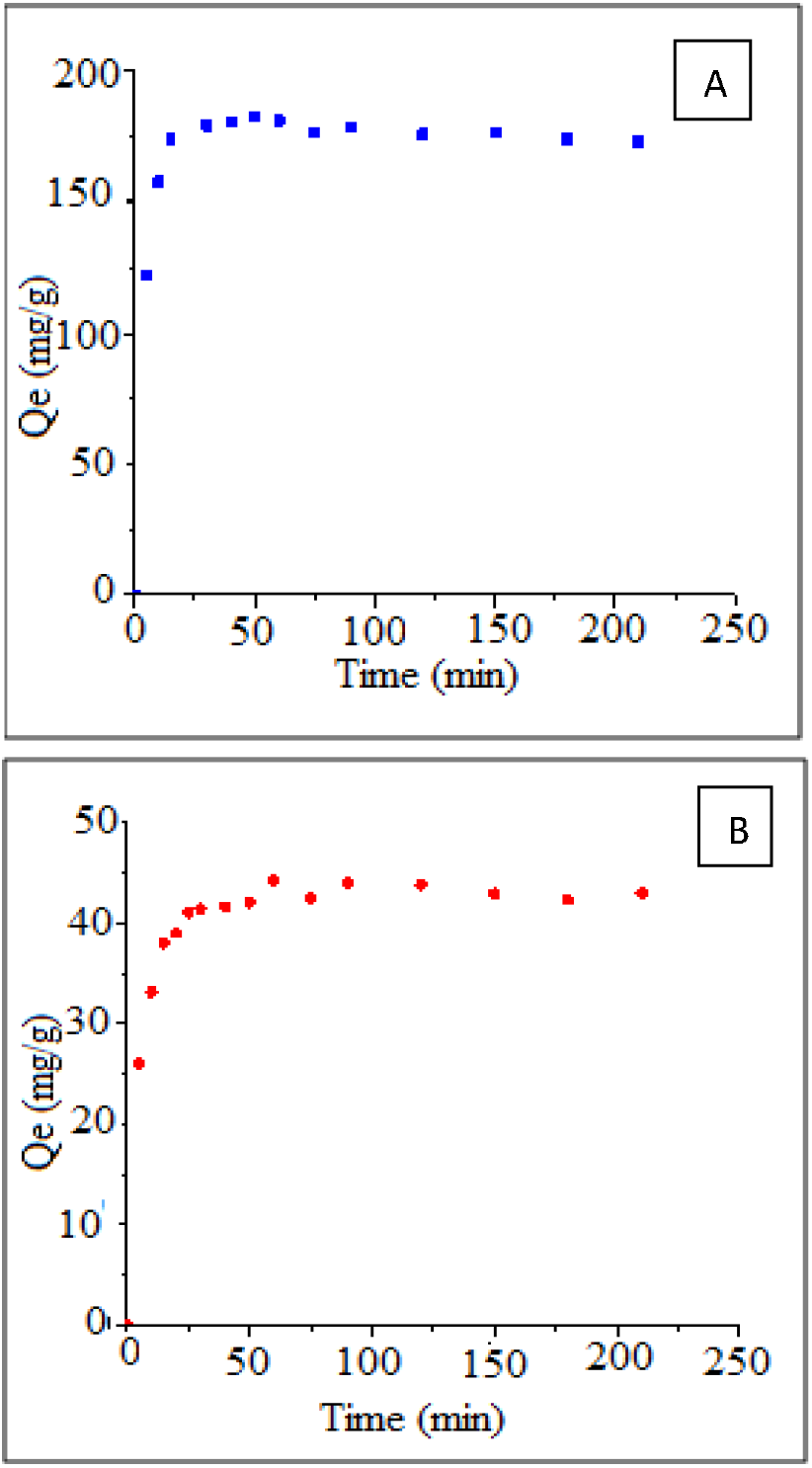
Kinetics of elimination of Janus GreenB by DMDDA-41C (A) and Orange II Sodium by DMDDA-41A (B)

In the two figures (A and B), two distinct phases of adsorption are observed: the first between 0 and 40 minutes which is rapid and takes place at the sites accessible on the surface, followed by a slow evolution of the adsorbed amount before to reach the second phase, which is in the form of an equilibrium plateau, due to the diffusion inside the pores of the material. The balance is reached after 60 minutes.

Figures 10 and 11 represent the linearization of the two kinetic models for the two organic pollutants and Table 8 groups together the values of the parameters of the two kinetic models applied to their elimination.

**Figure 10:**
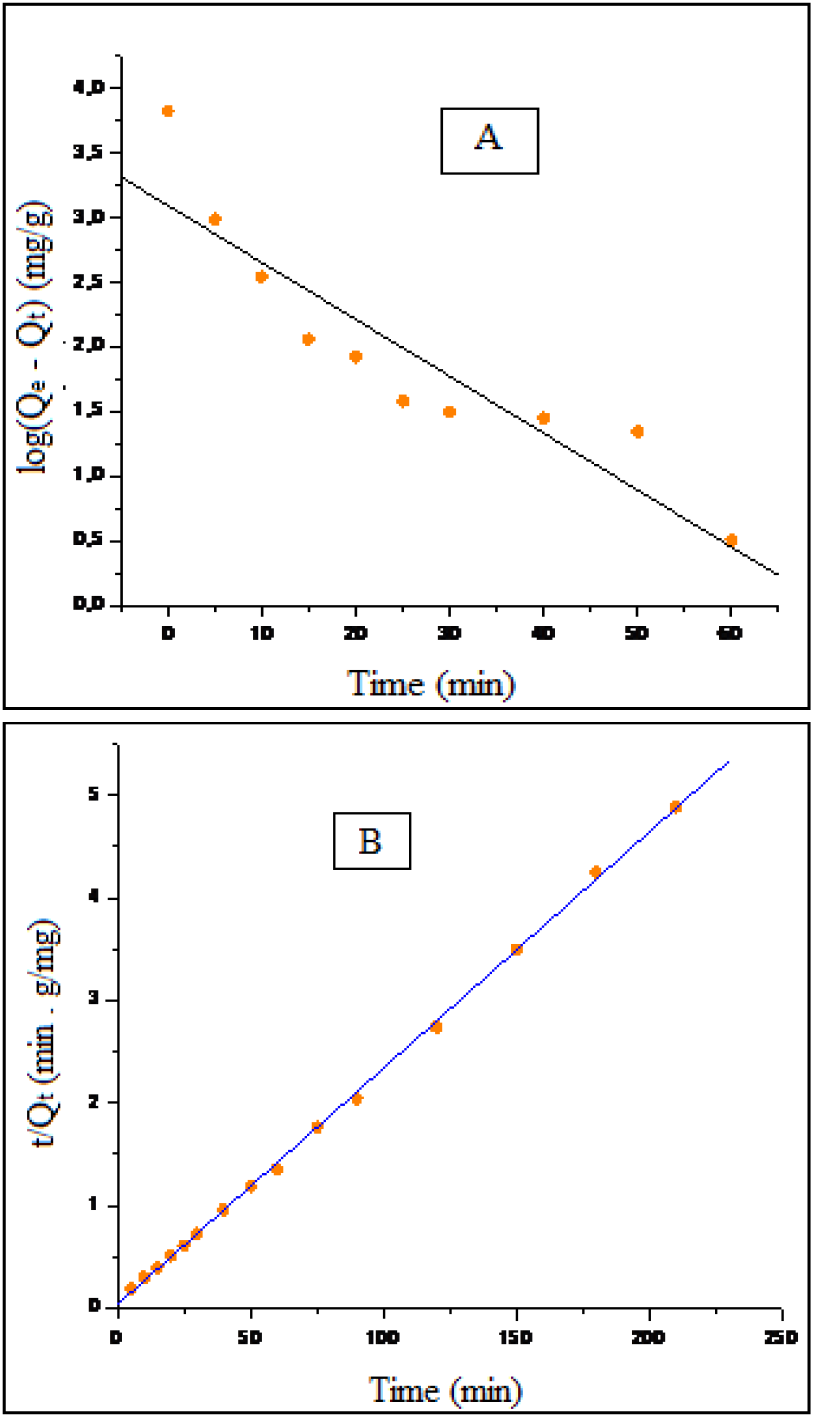
Application of the pseudo-first order (A) and pseudo-second order (B) model to the elimination of Orange II Sodium by DMDDA-41A.

**Figure 11:**
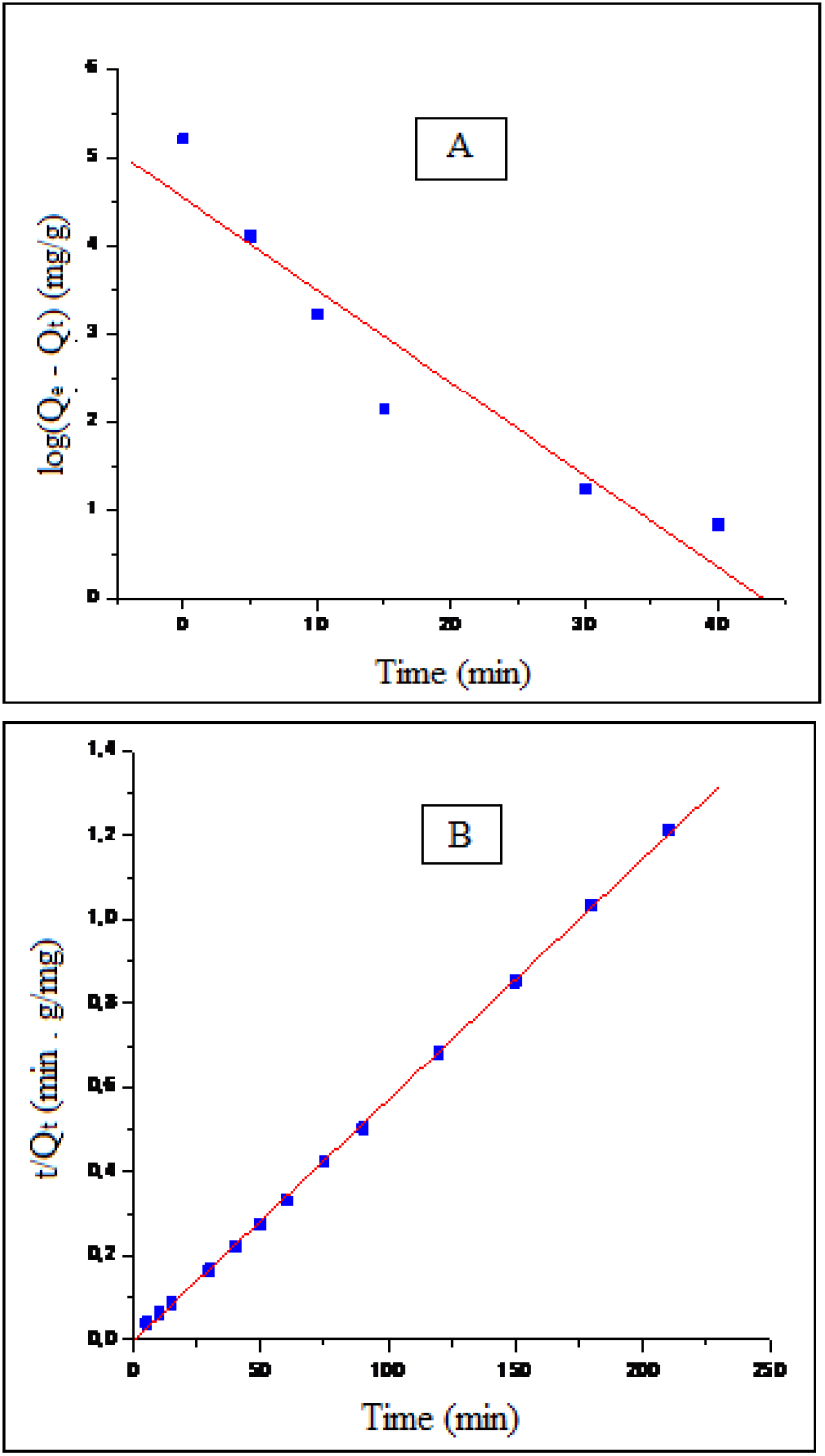
Application of the pseudo-first order (A) and pseudo-second order (B) model to the elimination of Janus Green B by the DMDDA-41C.

**Table 8:**
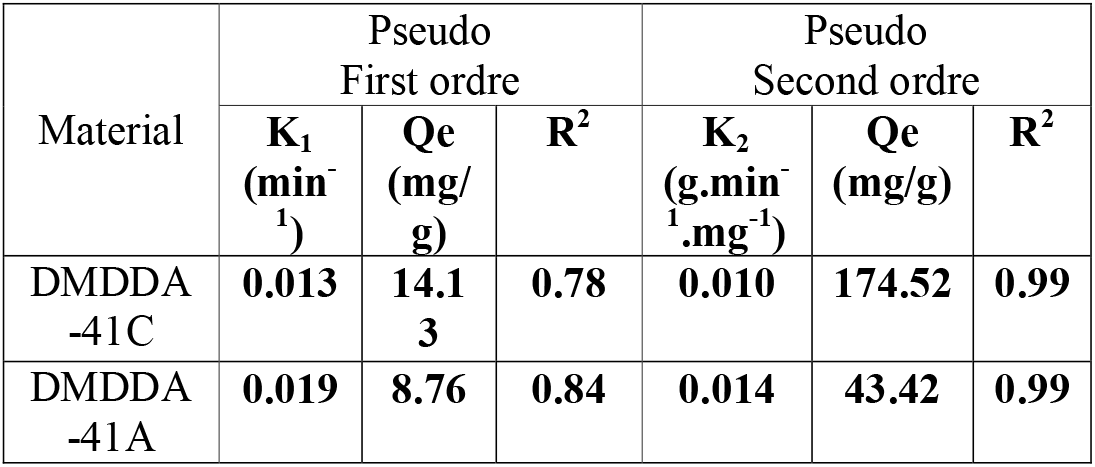
Parameters of the two kinetic models applied to the removal of the two organic pollutants "Janus Green B & Orange II Sodium".

| Material | Pseudo First ordre |  |  | Pseudo Second ordre |  |  |
| --- | --- | --- | --- | --- | --- | --- |
| | $K_1$<br>($\text{min}^{-1}$) | $Q_e$<br>($\text{mg/g}$) | $R^2$ | $K_2$<br>($\text{g} \cdot \text{min}^{-1} \cdot \text{mg}^{-1}$) | $Q_e$<br>($\text{mg/g}$) | $R^2$ |
| DMDDA-41C | **0.013** | **14.13** | **0.78** | **0.010** | **174.52** | **0.99** |
| DMDDA-41A | **0.019** | **8.76** | **0.84** | **0.014** | **43.42** | **0.99** |

According to the values of the kinetic constants, the kinetics of the dyes follows the pseudo second order model (R^2^= 0.999), rather than that of the pseudo-first.

The plots of the curves: Qt = f (t1 / 2), of the intraparticle diffusion (FIG. 12), are not linear and we notice three distinct sections associated with three successive steps:

- A first linear portion of the first step corresponds to a rapid diffusion in the boundary layer of the molecules of the solute, the adsorbate migrating from the solution to the external surface of the adsorbent.
- The second section, the curvature, is attributed to the intraparticle diffusion, which determines the rate of control of the adsorption mechanism.
- The plateau corresponds to a state of equilibrium: the intra-particle diffusion slows down, leading to a maximum of adsorption and a very low concentration of adsorbate in the solute [29]

**Figure 12:**
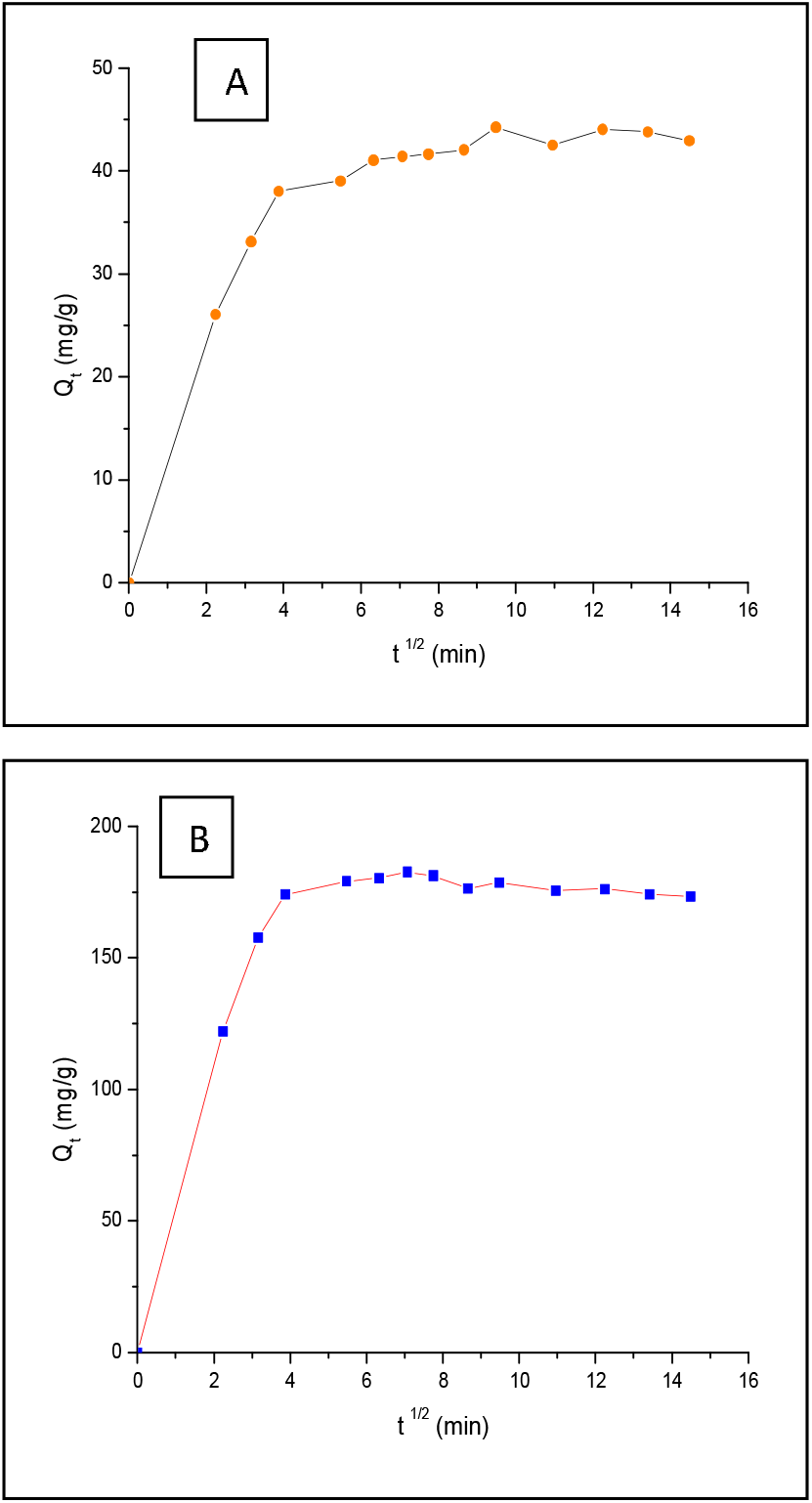
Intraparticular diffusion of Orange II Sodium (A) and Janus Green B (B)

**Table 9:**
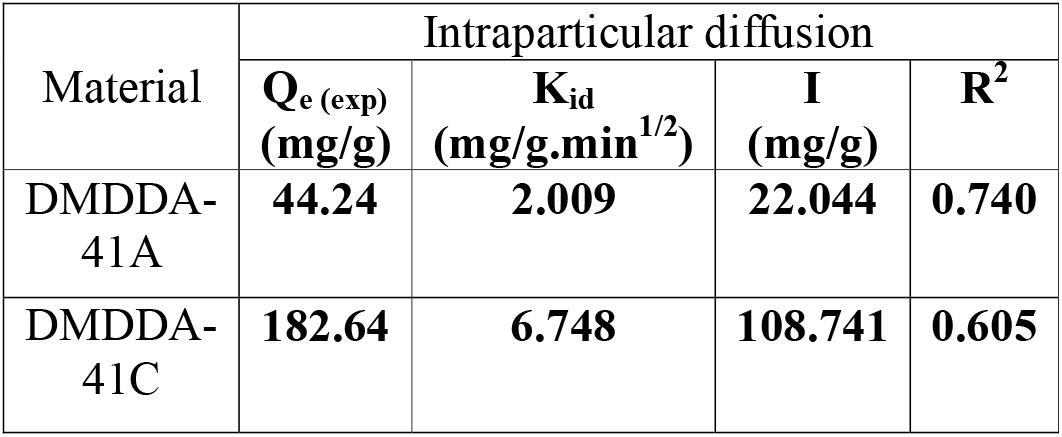
Kinetic parameters of the intraparticular diffusion of the two dyes "Janus Green B & Orange II Sodium".

The adsorption isotherms of the anionic and cationic dye at 25 ° C. by the materials DMDDA-41A and DMDDA-41C are shown in figure 13.

**Figure 13:**
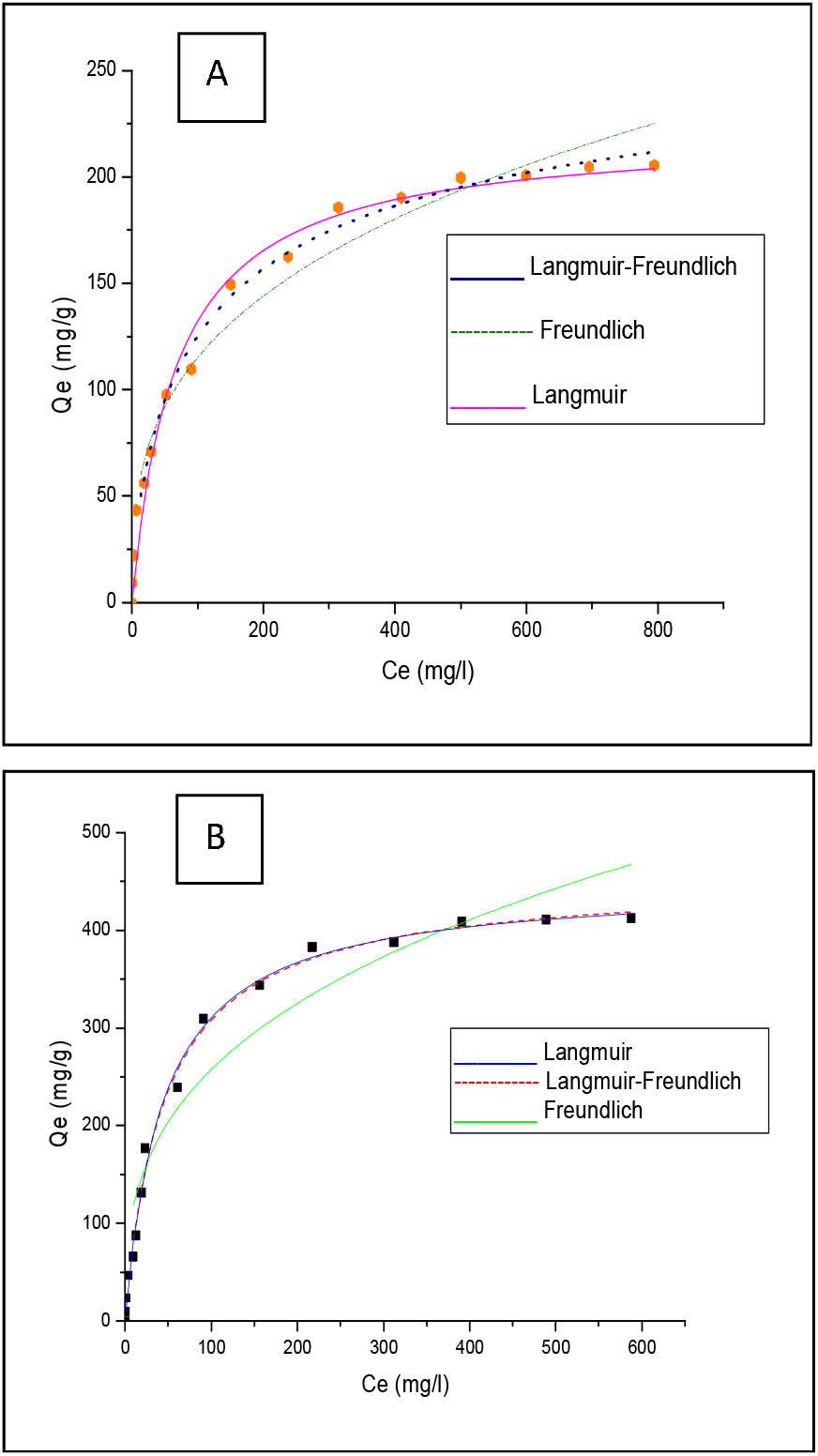
Adsorption isotherms of Orange II Sodium by DMDDA-41A (A) and Janus Green B by DMDDA-41C (B)

Using the classification of adsorption isotherms, those obtained are of type L "Langmuir". This type is characterized by a decreasing slope as the equilibrium concentration increases, due to the decrease in the number of adsorption sites, following the gradual covering of the surface of the various materials. In this type of adsorption, there is no interaction between the adsorbed molecules: this is a physisorption [28]. The representation of the adsorption isotherms is markedly better by the Sips model in the two dyes ***(R^2^> 0.994)*** and especially by the Langmuir model in the fixation of the cationic pollutant Janus Green B ***(R^2^> 0.995).***

The study of the linearizations of the Langmuir, Freundlich and Sips models (Langmuir-Freundlich), defined above, gives the various parameters of the three models that we have gathered in Table 10.

**Table 10:** Parameters of the adsorption isotherms applied to the removal of the two organic pollutants "Janus Green B & Orange II Sodium".

| Material | Langmuir |  |  | Freundlich |  |  | Sips (Langmuir-Freundlich) |  |  |  |
| --- | --- | --- | --- | --- | --- | --- | --- | --- | --- | --- |
| | $Q_{max}$ (mg/g) | $K_L$ (L/g) | $R^2$ | $K_F$ (L/g) | $1/n$ | $R^2$ | $Q_{max}$ (mg/g) | $K_S$ (L/g) | $n$ | $R^2$ |
| DMDDA-41A | 221.06 | 0.014 | 0.98 | 24.94 | 0.323 | 0.97 | 178.02 | 0.04 | 0.65 | 0.99 |
| DMDDA-41C | 448.00 | 0.022 | 0.995 | 54.77 | 0.336 | 0.94 | 455.23 | 0.02 | 0.95 | 0.99 |

The essential characteristic of the Langmuir isotherm can be expressed as a function of a dimensionless constant separation factor which is given by the following expression:

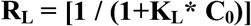

The evolution of the separation factors as a function of the initial concentration at 25 ° C is shown in figure 14. For both materials and the two dyes, the values of this factor are between ***0 <RL <1***: This shows that the adsorption of the two dyes by the different materials is a favorable process, and the ***1 / n*** values indicate a favorable adsorption especially for the low concentrations [26, 27].

## 5. Conclusion

The main objective of this work was to synthesize the MCM-41 material from optimized protocols. It was then possible to increase the size of the pores and the specific surface area of the parent material by the post-synthesis incorporation of blowing agents (long chain carbon amines DMDDA).

The results of X-ray diffraction characterization of the different matrices show that for the five types of morphology the diffractogram obtained has three indexable peaks in (100), (110) and (200) in hexagonal symmetry (space group P6mm). For amine and calcined materials, the presence of indexable peaks in hexagonal network always shows that the structure is always preserved after the post-synthesis addition of the amine emulsion or by a high treatment temperature.

The BET analysis of the materials shows a remarkable decrease in pore diameter and the specific surface area for each of the amines and deaminates, unlike the MCM-41C which has the highest surface area and pore diameter.

The thermogravimetric analysis shows a thermal stability of the materials with a greater loss of mass for the amine material, which again confirms the incorporation of the amines inside the pores.

The results of the infrared spectra show the incorporation of the NH_2_ groups and for those of the zeta-metry show that the surface charge differs from one material to another, which suggests their applications in the adsorption domain of the various micropollutants.

The results obtained during the adsorption study show the effectiveness of these materials, in particular "the amine material" and "the calcined material" for decolorizing aqueous media contaminated with organic dyes. The kinetic studies and the adsorption isotherms were carried out to clarify the method of fixing each of the two dyes on the two materials tested. The experiments showed that the amine material had a maximum capacity for fixing Orange II (221.06 mg / g) (anionic dye), whereas for a maximum adsorption capacity (455.23 mg / g) of Janus GB (cationic dye), the calcined material is more efficient.

The second order kinetic model applies well in the case of the adsorbent / adsorbate systems studied (R^2^ = 0.999). The adsorption isotherms of the two dyes by the two adsorbents are satisfactorily described by the Langmuir model in the case of fixation of the cationic pollutant (R^2^ = 0.995). However, for the adsorption of the anionic pollutant, the Sips model is more favorable (R^2^ = 0.994).

The RL separation factor values for the Langmuir model of the two dyes is between (0 <RL <1): this shows that the adsorption in this case is a favorable process.

The adsorption of the two micropollutants (Janus Green B and Orange II Sodium) is favored at low concentrations (1 / n <1).

